# Normal Somitogenesis Despite the Absence of *Snai1* in the Presomitic Mesoderm

**DOI:** 10.64898/2026.10.27.762210

**Authors:** Ana Casaca, Rita Aires, Ana Nóvoa, Moisés Mallo

## Abstract

During vertebrate somitogenesis, *Snai1* is expressed dynamically in the presomitic mesoderm (PSM), oscillating in phase with core segmentation clock genes. Whether this reflects a functional requirement for *Snai1* in clock activity has remained unresolved, largely because global or epiblast-specific loss of *Snai1* causes early embryonic lethality. Here, we identify a conserved enhancer downstream of the *Snai1* locus (SCDE) that recapitulates major domains of endogenous *Snai1* expression, including the PSM, tailbud, cranial mesenchyme, branchial arches and forelimb buds. Functional dissection identifies two domains within this enhancer, B1 and B3, as partially redundant core regulatory elements that together account for most SCDE activity in the posterior mesoderm. Deletion of these modules strongly reduces *Snai1* expression in the PSM while preserving its dynamic expression pattern and *Lfng* oscillations, without producing any detectable skeletal phenotype. To directly assess the requirement for *Snai1*, we generated conditional mutants lacking *Snai1* throughout the posterior mesoderm, including the PSM. Despite the total absence of *Snai1* expression in this area, these embryos show normal somite formation, compartmentalisation and segmentation clock activity. Unexpectedly, complete loss of *Snai1* in this territory results in a fully penetrant gain of one pre-sacral vertebra, pointing to a role in fine-tuning axial segment number rather than in segmentation itself. Together, our findings identify SCDE as a bona-fide cis-regulatory element involved in the control of *Snai1* expression in the PSM, uncouple its oscillatory expression from an essential role in the segmentation clock and suggest that this gene is dispensable for somitogenesis.

## 1. Introduction

During vertebrate embryogenesis, precise spatial and temporal control of gene expression is essential for the coordinated formation of body structures. One key process exemplifying this regulation is somitogenesis, the periodic segmentation of the paraxial mesoderm into somites that give rise to the vertebral column [1]. Segmentation occurs at the anterior end of the presomitic mesoderm (PSM), the unsegmented tissue spanning from the posterior growth zone, where new mesodermal cells are continuously added, to the most recently formed somite. This process is controlled by a molecular oscillator, the segmentation clock, together with signalling gradients [2] that confer positional identity along the posterior-to-anterior extent of the PSM, defining where posterior border of each new somite will form. Together, these mechanisms ensure the rhythmic and sequential formation of somites from the posterior growth zone [3].

Members of the *Snail* gene family are among the key transcription factors implicated in mesodermal differentiation [4]. *Snai1* encodes a zinc-finger transcriptional repressor best known for its role in epithelial–mesenchymal transition, cell migration, and tissue remodelling [5,6]. In the mouse embryo, *Snai1* shows a dynamic expression in various embryonic domains, including the primitive streak, posterior mesoderm, PSM, cranial mesenchyme, and branchial arches [7–10].

Within the PSM, *Snai1* expression has been reported to oscillate in phase with other segmentation clock genes, suggesting its participation in clock function [11,12]. However, functional analysis of *Snai1* in this tissue has been hindered by the early developmental arrest when globally inactivated or when conditionally removed from the epiblast [7,13]. Whether oscillatory expression in the PSM reflects an essential requirement for *Snai1* in the segmentation clock therefore remains unresolved. Addressing this question requires tools capable of manipulating *Snai1* activity specifically within the PSM, without the confounding early lethality associated with global or epiblast-specific loss. Thus, a closer examination of the *cis*-regulatory elements that control its tissue-specific expression is important.

The transcriptional regulation of *Snai1* mirrors the complexity of its developmental functions. Several upstream pathways, including those from the TGFB, WNT and NOTCH signalling families, modulate *Snai1* transcription [5,6,14]; as reviewed in [4,15], yet large portions of its genomic locus remain uncharacterized. Increasing evidence points to the contribution of various *cis*-regulatory elements, illustrated by the recent identification of long-range regulatory element at the *Snai1* locus in cancer cells [16]. Most studies of *Snai1* regulatory elements, however, have been conducted in cancer or cell line models, with very few addressing its transcriptional regulation in the embryonic context. One exception is an element located downstream of the *Snai1* locus, first identified in melanoma cells [17,18], which was later shown to be a direct transcriptional target of Msgn1, a master regulator of paraxial mesoderm differentiation, controlling *Snai1* expression in the PSM [19]. However, its internal regulatory architecture, full enhancer activity across embryonic territories and functional contribution to *Snai1* expression and PSM biology remain poorly characterised.

Here, we further characterise the *Snai1* downstream regulatory region and show that it functions as a modular enhancer controlling *Snai1* expression in the PSM and other embryonic territories. Mice lacking this enhancer exhibit a selective reduction of *Snai1* in the PSM, yet display normal segmentation and somitogenesis, indicating that *Snai1* activity in this tissue is largely dispensable for somite formation. This conclusion is further supported by the conditional inactivation of *Snai1* in the PSM and posterior mesoderm, which similarly leaves somitogenesis intact. Together, these findings indicate that, contrary to assumptions based on its oscillatory expression, *Snai1* is not required for the segmentation process itself. Notably, conditional inactivation in the posterior mesoderm resulted in a consistent gain of one pre-sacral vertebra, suggesting a role for *Snai1* in fine-tuning axial segment number rather than in segmentation *per se*. This work thus reveals an uncoupling between the oscillatory transcriptional behaviour of *Snai1* and its actual functional requirement during vertebrate axis formation.

## 2. Materials and methods

### 2.1 Ethics statement

All animal procedures were performed in accordance with Portuguese (Portaria 1005/92) and European (Directive 2010/63/EU) legislation and guidance on animal use in bioscience research. The project was reviewed and approved by the Ethics Committee the Gulbenkian Institute for Molecular Medicine (GIMM) and by the Portuguese National Entity ‘Direcção Geral de Alimentação Veterinária’ (license reference: 014308).”

### 2.2 Mouse strains and genotyping

The *Snai1* floxed mouse line (*Snai1*^flox/flox^) [20] was obtained from The Jackson Laboratory (B6.129S-*Snai1*^tm2Grid^/J; Stock No. 010686). A constitutive *Snai1* null allele (*Snai1*^−^) was generated by crossing *Snai1*^flox/flox^ mice with the Meox2-Cre germline deleter strain (B6.129S4-*Meox2*^tm1(cre)Sor^/J; Stock No. 003755), as previously described [21]. Conditional *Snai1* mutant embryos (*T*-Cre^+/0^:*Snai1*^flox/−^, hereafter *cSnai1*) were generated by crossing *T*-Cre^+/0^:*Snai1*^+/−^ males with *Snai1*^flox/flox^ females. In experiments involving *Snai1* inactivation, control embryos and pups were littermates of genotypes containing at least one functional *Snai1* allele in *trans* (including *Snai1^flox/flox^*, *Snai1^flox/−^*, and *T-Cre^+/0^:Snai1^flox/+^*), none of which showed a phenotype distinguishable from wild-type.

The ROSA26-lacZ-R [22] line was used to assess *T-Cre* recombination activity by crossing *T-Cre^+/0^*males with *ROSA26-lacZ-R* females and recovering embryos at different embryonic stages for β-galactosidase staining.

Genotyping was performed by PCR using genomic DNA extracted from ear or digit biopsies for adult mice, yolk sacs for embryos up to E11.5, or intestinal tissue for later staged fetuses (E17.5–E18.5). For adult tissue, samples were incubated in 50 μL of PBND buffer (50 mM KCl, 10 mM Tris-HCl pH 8.3, 2.5 mM MgCl2, 0.1 mg/mL gelatin, 0.45% NP40, 0.45% Tween-20) supplemented with 100 μg/mL proteinase K at 55°C overnight, followed by heat-inactivation at 95°C for 15 minutes. For yolk sac DNA extraction, tissue was incubated in 50 μL of lysis buffer (50 mM KCl, 10 mM Tris-HCl pH 8.3, 2 mM MgCl2, 0.45% NP40, 0.45% Tween-20) supplemented with 100 μg/mL proteinase K at 55°C overnight, followed by heat-inactivation at 95°C for 15 minutes. For intestine samples, tissue was incubated in 500 ul LAIRDS buffer supplemented with 100 μg/mL proteinase K at 55°C overnight, then precipitated with 500 ul Isopropanol and resuspended in 250 μL TE. 1 μL of genomic DNA was used per PCR reaction. Primer sequences and expected product sizes are listed in Supplementary Table1.

### 2.3 Generation of transgenic embryos and lines

The SCDE region was amplified by PCR from genomic DNA (primers in Supplementary Table1) and cloned into a vector containing the adenovirus major late promoter, the β-galactosidase coding region and the SV40 polyadenylation signal [23]. Constructs carrying individual blocks (B1, B2, B3) were generated by standard PCR amplification from this template (primers in Supplementary Table1). Block-deletion constructs (ΔB1, ΔB2, ΔB3) were generated using an overlapping PCR mutagenesis strategy (primers in Supplementary Table1). The ΔB1B3 construct was generated by replacing the XhoI–BglII fragment of the ΔB1 construct with the corresponding XhoI–BglII fragment from the ΔB3 construct. All constructs were verified by direct sequencing.

To generate the T-Cre transgenic line the cre recombinase was cloned downstream of the *Tbxt* primitive streak-specific promoter [24] and upstream of the SV40 polyadenylation signal. For microinjection, constructs were isolated from the plasmid backbone by restriction digestion, gel-purified using the NZYGelpure kit (NZYTech #MB01102) and eluted in 50 μL of the kit’s elution buffer. Purified DNA was diluted to 2 ng/μL in microinjection buffer (10 mM Tris-HCl, 0.25 mM EDTA, pH 7.5) and microinjected into the pronuclei of fertilized FVB/N oocytes according to standard protocols [25]. Microinjected oocytes were transferred into the uteri of pseudopregnant NMRI females and embryos were recovered at different stages for β-galactosidase staining. To establish the T-Cre line, founder mice carrying the transgene were identified by PCR genotyping and backcrossed to generate the stable line.

### 2.4 Generation of CRISPR/Cas9 mouse lines

The ΔB1B3 mouse line was generated by CRISPR/Cas9-mediated genome editing using the Cas9-mSA/biotinylated donor strategy as previously described [26]. Two guide RNAs flanking Blocks 1 and 3 (Supplementary Table1) were co-injected with Cas9-mSA mRNA and a biotinylated donor ssDNA into fertilized FVB/N oocytes. The donor was generated by PCR amplification of the ΔB1B3 reporter construct using a biotinylated forward primer (Supplementary Table1) and contained the endogenous *Snai1* enhancer sequence lacking Blocks 1 and 3, flanked by 200 nt homology arms matching the sequences immediately flanking the two target sites.

The ΔSCDE line was generated by CRISPR/Cas9-mediated deletion as previously described [27]. Two guide RNAs targeting sequences flanking the SCDE region (sequences in Supplementary Table1) were co-injected with Cas9 mRNA and a synthetic ssDNA donor containing 120 nt, comprising 60 nt homology arms matching the sequences immediately flanking the deletion breakpoint, with no additional sequence between them, resulting in a precise deletion of the intervening SCDE sequence.

For both CRISPR-generated lines, oocyte transfer was performed as described above. Founder animals were screened by PCR genotyping and the targeted alleles were confirmed by direct sequencing.

### 2.5 β-galactosidase staining

Embryos were dissected in ice-cold PBS and fixed in 4% paraformaldehyde at 4°C for 30 minutes. Following three washes in β-gal wash buffer (PBS containing 0.02% Tween-20) for 10 minutes at room temperature, β-galactosidase activity was developed overnight at 37°C in the dark in PBS containing 5 mM K_3_Fe(CN)_6_, 5 mM K_4_Fe(CN)_6_, 2 mM MgCl_2_, 0.02% Tween-20 and 0.4 mg/mL X-gal (Promega #V3941). The reaction was stopped with β-gal wash buffer, embryos were post-fixed in 4% paraformaldehyde overnight at room temperature and stored in PBS. For vibratome sectioning, post-stained embryos were embedded in a mixture of 0.45% gelatin, 27% bovine serum albumin and 18% sucrose in PBS, jellified with 1.75% glutaraldehyde, and sectioned at 20 μm.

### 2.6 Whole-mount in situ hybridisation

Whole-mount *in situ* hybridisation was performed as previously described [28]. A digoxigenin-labelled antisense Snai1 RNA probe (795 bp) corresponding to the complete coding sequence was generated by cloning a PCR-amplified fragment from pCDNA3-mSnai1 (primers in Supplementary Table1) into a vector containing a T3 promoter, followed by in vitro transcription with T3 RNA polymerase in the presence of DIG-UTP according to the manufacturer’s instructions. Embryos were recovered at different embryonic stages, fixed in 4% paraformaldehyde overnight at 4°C and processed according to standard protocols. All staining experiments included mutant and wild-type littermate embryos processed in parallel under identical conditions. The number of embryos examined per probe, genotype and stage is indicated in the corresponding figure legends and text, and varied according to litter size and embryo availability. In all cases, staining patterns were consistent across replicate embryos of the same genotype and stage, and representative images are shown.

### 2.7 Skeletal preparation and analysis

Foetuses were recovered by caesarean section at E18.5 and skeletal phenotypes analysed by Alcian Blue/Alizarin Red staining as previously described [28]. Vertebral formulae were determined by examination of cleared and stained skeletal preparations and scored according to standard anatomical criteria.

### 2.8 Quantification and statistical analysis

Phenotypic data from transgenic reporter assays, enhancer deletion lines and conditional mutants are reported as raw counts and proportions of embryos or animals displaying a given expression pattern, phenotype or vertebral formula, scored by direct visual assessment of β-galactosidase staining, *in situ* hybridisation signal or skeletal preparations. Given the categorical, presence/absence nature of these scores and the expected biological variability between independent transgenic insertions, no formal statistical tests were applied to compare proportions between genotypes or lines; results are presented as fractions throughout the text and figures.

## 3. Results

### 3.1 A conserved downstream enhancer controls axial and craniofacial Snai1 expression

Previous work identified a region 3′ to the *Snai1* transcriptional unit as a target of Msgn1 in the PSM [19]. To dissect the regulatory potential of this locus, we performed a comparative genomic analysis and identified a broader conserved region spanning +4540 to +5955 bp downstream of the *Snai1* transcriptional start site (*Snai1* Conserved Downstream Enhancer - SCDE; Fig. 1A), which includes and extends beyond the previously characterised Msgn1-responsive sequences (Supplementary Fig. 1). To test whether this region functions as a developmental enhancer, we generated transgenic mice carrying a SCDE-lacZ reporter and assessed its activity between E8.5 and E10.5, stages at which *Snai1* is expressed in the primitive streak, nascent mesoderm, cranial mesenchyme, branchial arches, PSM, somites and posterior tailbud, with forelimb bud expression becoming evident from E9.5 onwards [8–10] (Fig. 1B, a-c). SCDE-lacZ reporter embryos showed β-galactosidase activity that largely mirrored the endogenous *Snai1* expression pattern across all stages examined (Fig. 1B, compare a-c with d-f), with the PSM, somites and posterior tailbud showing the most robust and consistent activity across embryos. Craniofacial expression, encompassing head mesenchyme and branchial arches, was similarly observed at all stages examined (Fig. 1B, d-f). Forelimb bud activity emerged from E9.5 (Fig. 1B, e, red arrowhead) and was maintained at E10.5 (Fig. 1B, f) consistent with the onset of endogenous *Snai1* expression in this domain. These results establish SCDE as a bona fide developmental enhancer whose activity recapitulates the major domains of *Snai1* expression *in vivo*. To provide a quantitative characterisation of this activity, we analysed a total of 9 SCDE-lacZ transgenic embryos at E10. Of these, two displayed ubiquitous staining inconsistent with endogenous *Snai1* expression, likely due to position effects associated with the integration site and were excluded from further analysis. The remaining 7 embryos showed domain-specific β-galactosidase activity in the PSM and somites (6/7), posterior tailbud (7/7), head mesenchyme (7/7), branchial arches (6/7) and forelimb bud staining (6/7; Fig. 2A, SCDE; representative embryo in Fig. 2B, a, f). It should be noted that, while β-galactosidase staining is observed in the PSM, the high β-galactosidase stability does not allow capturing the oscillatory activity that characterises *Snai1* expression.

**Figure 1.**
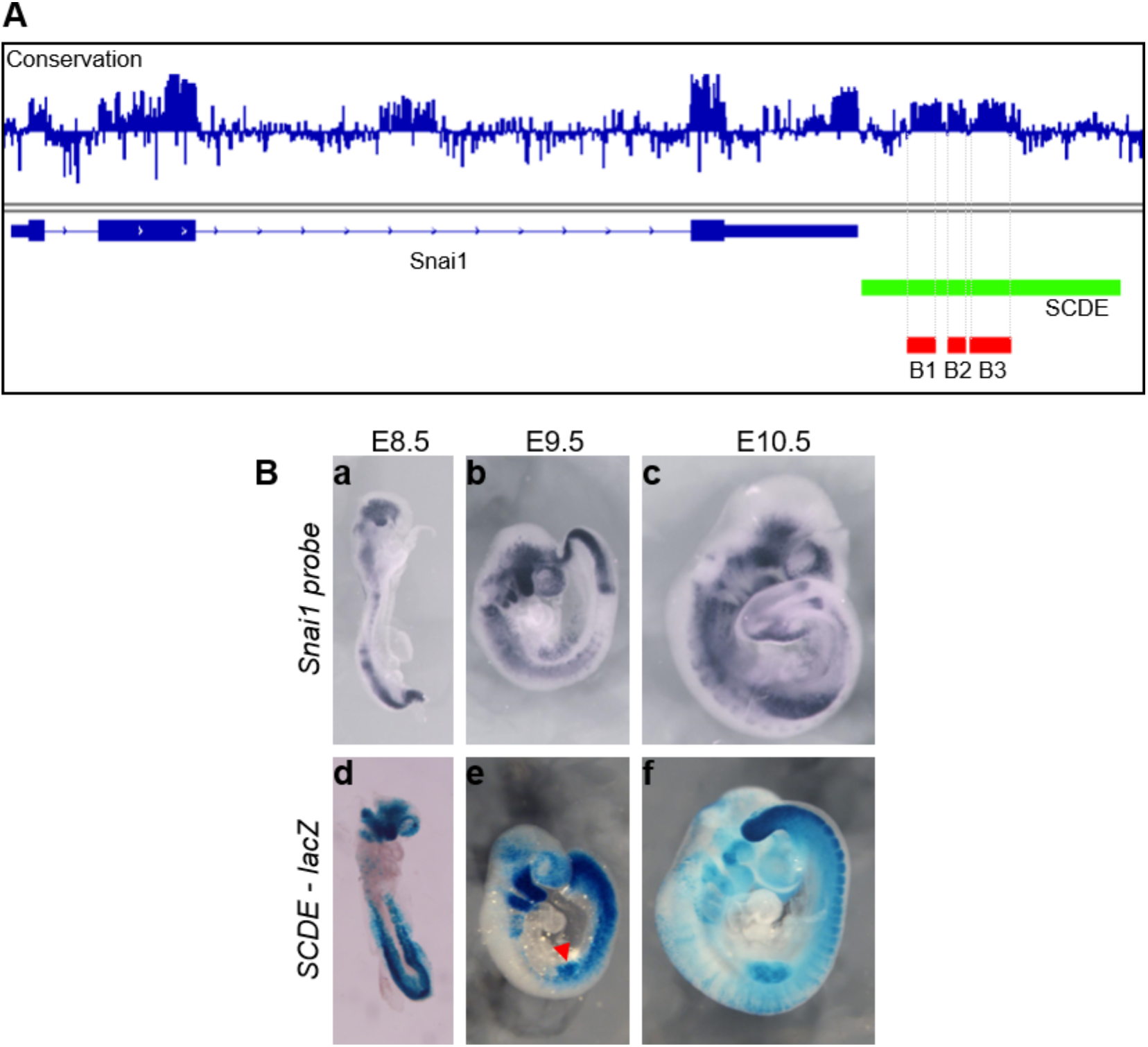
A conserved downstream enhancer recapitulates major domains of *Snai1* expression. **A:** Schematic representation of the *Snai1* locus showing sequence conservation across vertebrates (top track), gene structure with exons (dark blue boxes) and introns (arrows indicating direction of transcription), and the downstream SCDE region (green), highlighting the three conserved blocks, B1, B2 and B3 (red). **B:** Whole-mount *in situ* hybridisation for endogenous *Snai1* (a–c) compared to whole-mount β-galactosidase staining of SCDE-lacZ transgenic reporter embryos (d–f) at E8.5 (a, d), E9.5 (b, e) and E10.5 (c, f). SCDE-lacZ activity recapitulates major domains of endogenous *Snai1* expression, including the presomitic mesoderm (PSM), somites, posterior tailbud, craniofacial mesenchyme and limb buds. The red arrowhead in (e) indicates forelimb bud activity at E9.5.

**Figure 2.**
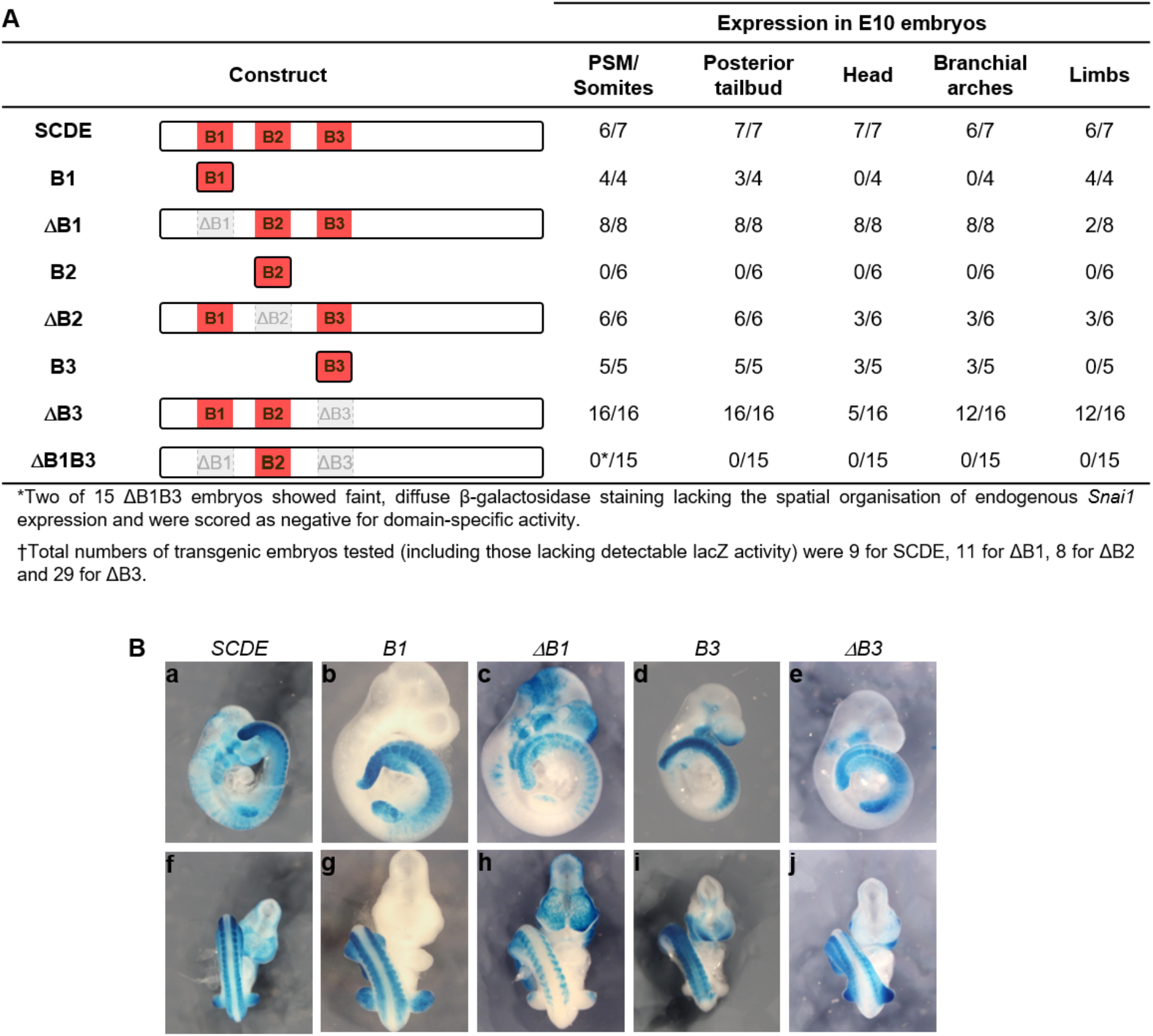
Modular dissection of the SCDE enhancer identifies B1 and B3 as partially redundant core regulatory elements. **A:** Schematic of the SCDE reporter constructs tested, including the full-length enhancer, individual conserved blocks (B1, B2 and B3), and deletion constructs (ΔB1, ΔB2, ΔB3 and ΔB1B3). The frequency of E10 transgenic embryos displaying β-galactosidase activity in each anatomical domain is shown for each construct. For SCDE, ΔB1, ΔB2 and ΔB3, a subset of transgenic embryos showed no detectable β-galactosidase activity, likely reflecting positional effects associated with the transgene insertion site; these embryos were excluded from domain-specific scoring, and fractions represent the proportion of embryos with detectable reporter activity that displayed expression in each domain (†). **B:** Representative whole-mount β-galactosidase-stained embryos for SCDE, B1, ΔB1, B3 and ΔB3 constructs, shown in lateral (a–e) and dorsal (f–j) views. The analysed domains include PSM/somites, posterior tailbud, head mesenchyme, branchial arches and limb buds.

To map the functional sub-regions of the SCDE, we subdivided it into three blocks based on their sequence conservation across vertebrates (B1, B2 and B3; Fig. 1A and 2A) and assessed their activity focusing on E10 embryos. B1 alone drove β-galactosidase expression in the PSM and somites (4/4), posterior tailbud (3/4) and forelimb buds (4/4; Fig. 2A; Fig. 2B, b, g). In contrast, head mesenchyme and branchial arch expression were absent in all four B1 embryos, indicating that B1 reproduces the axial and limb components of the full SCDE pattern but is insufficient to drive anterior craniofacial activity. To directly test the contribution of B1 to the SCDE activity, we analysed embryos carrying a SCDE construct lacking B1 (ΔB1). Eight out of 11 transgenic embryos showed β-galactosidase staining. They all preserved axial expression in the PSM and somites, posterior tailbud, head mesenchyme and branchial arches (Fig. 2A; Fig. 2B, c, h), demonstrating that this region is not necessary for SCDE activity and suggesting the existence of additional regions that can act redundantly with B1 in these regions. Forelimb bud staining was, however, markedly reduced, detected in only 2/8 ΔB1 embryos compared to 4/4 with B1 alone and 6/7 with the full SCDE. These observations identified B1 as the primary driver of limb enhancer activity within the SCDE. Notably, head mesenchyme and branchial arch staining was consistently more prominent in ΔB1 embryos than in full SCDE controls (compare Fig. 2B, c, h with a, f), raising the possibility that B1 might restrain enhancer activity in craniofacial domains, perhaps through competition or interference of adjacent regulatory sequences.

To test whether this modularity extends to other conserved blocks, we analysed B3, another of the highly conserved regions within SCDE. B3 alone drove β-galactosidase expression in the PSM, somites and posterior tailbud in all 5 transgenic embryos analysed and in head mesenchyme and branchial arches in 3/5 embryos (Fig. 2A; Fig. 2B, d, i). Forelimb bud expression was, however, absent in all 5 B3 embryos, indicating that B3 lacks regulatory activity in this domain. The contrasting forelimb phenotypes of B1 (4/4) and B3 (0/5) therefore identify B1 as the sole sub-element of SCDE with limb enhancer activity. To further assess B3 contribution to the axial and craniofacial SCDE activities, we analysed embryos carrying a B3 deletion (ΔB3). From 29 transgenic embryos 16 showed β-galactosidase staining. They all retained full axial and posterior expression in the PSM, somites and tailbud (16/16; Fig. 2A; Fig. 2B, e, j). Head mesenchyme staining was observed in 5/16 embryos, whereas branchial arch staining was detected in 12/16, indicating that these two craniofacial domains are differentially affected by the loss of B3. These results, together with those involving B1, suggest that B1 and B3 might act redundantly to control expression in the PSM, somites, tailbud and, to a slightly reduced level also in the branchial arches. However, SCDE head mesenchyme activity seems to be more dependent on B3.

To directly test the potential B1 and B3 redundant activity in axial and posterior regions of the embryo, we generated a construct lacking both blocks (ΔB1B3). 13 out of 15 transgenic embryos completely lacked β-galactosidase activity and the remaining two showed faint, diffuse staining lacking the spatial organisation of endogenous *Snai1* expression, consistent with background (Fig. 2A). The near-complete loss of enhancer activity upon simultaneous removal of B1 and B3, confirms the redundant and specific roles of these blocks, and establish them as the core functional modules of the SCDE.

These data also suggest that the other conserved region within SCDE, namely B2, might not be involved in the activity of the enhancer. The absence of detectable β-galactosidase expression in any of the 6 transgenic embryos in which the B2 region was tested alone and the fairly normal activity of SCDE lacking the B2 region (ΔB2) in 6 out of 8 embryos showing staining in the PSM and somites (6/6), posterior tailbud (6/6), head mesenchyme (3/6), branchial arches (3/6) and forelimb buds (3/6; Fig. 2A) further support that B2 lacks both autonomous enhancer activity and any significant modulatory role within the intact SCDE.

### 3.2 Deletion of the Downstream Enhancer Selectively Reduces Snai1 in the Presomitic Mesoderm

To directly assess the contribution of the SCDE to endogenous *Snai1* regulation, we generated mouse lines carrying targeted deletions using CRISPR/Cas9 genome editing. Based on the enhancer dissection using the transgenic reporters, two deletion strategies were implemented: combined removal of Blocks 1 and 3 (ΔB1B3), which encompass the core enhancer modules driving axial expression, and deletion of the entire conserved region (ΔSCDE; Supplementary Fig. 2A). Homozygous mutants for both deletions were viable, fertile and showed no gross morphological abnormalities during embryogenesis or postnatally. Skeletal analysis further revealed a normal vertebral formula (C7T13L6S4) in all ΔB1B3 and ΔSCDE homozygous mutants examined (n=8 for each genotype; data not shown).

To evaluate the contribution of SCDE to *Snai1* transcription, we performed whole-mount *in situ* hybridisation on ΔB1B3 homozygous mutants and wild-type littermates processed in the same staining experiment at E8.5, E9.5 and E10.5 (Fig. 3A-C). At E8.5, expression in non-PSM domains including the cranial mesenchyme, epiblast and nascent mesoderm was indistinguishable from wild-type controls in both intensity and spatial distribution, indicating that early *Snai1* expression domains were unaffected by the deletion (Fig. 3A, a, b). In the PSM, wild-type embryos at this stage displayed a characteristic pattern comprising two spatially distinct components: a broad, intensely stained posterior domain (Fig. 3A, red lines in g-i) and one, two or three progressively fainter anterior stripes (Fig. 3A, orange arrowheads in g-i, m-o). Notably, unlike the dynamic oscillatory expression previously described [11], this pattern was largely consistent across the 9 wild-type embryos analysed, suggesting that at E8.5 *Snai1* is expressed in the PSM following a distinct pattern rather than the oscillatory behaviour characteristic of the mature segmentation clock. In ΔB1B3 mutants, the broad posterior domain was markedly reduced or absent (thinner red lines in Fig. 3A, j-), while the anterior stripes were comparatively less affected (Fig. 3A, j–l, p–r; n=10), indicating that SCDE selectively regulates the posterior PSM component at this stage.

**Figure 3.**
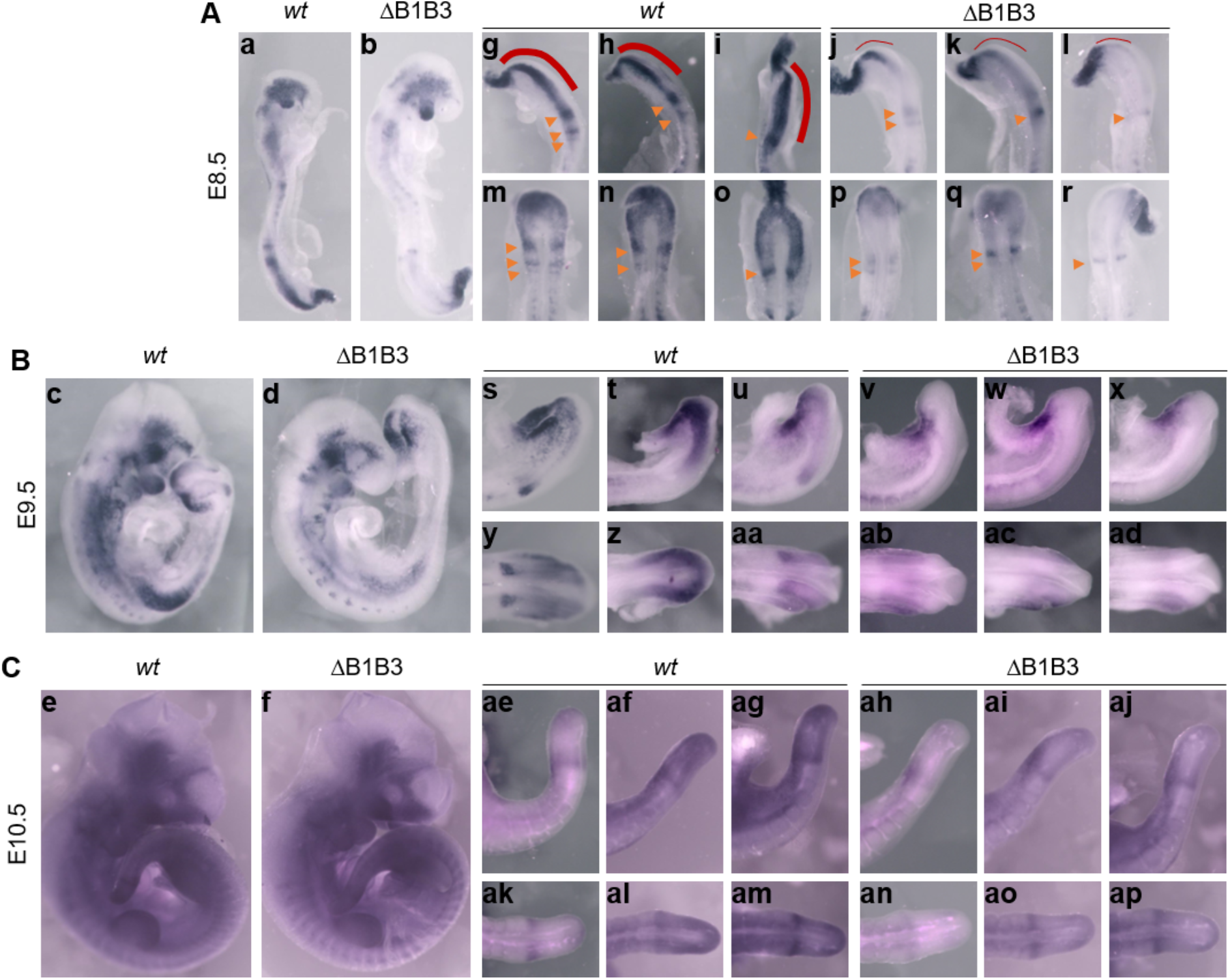
Deletion of B1 and B3 strongly reduces *Snai1* expression in the PSM while preserving its dynamic pattern. Whole-mount *in situ* hybridisation for Snai1 in *wild-type* littermates and ΔB1B3 embryos. **A:** E8.5. Whole embryos (a, b) and higher-magnification lateral (g–l) and dorsal (m–r) views of the posterior PSM/tailbud. Red lines indicate the broad posterior expression domain (thicker lines: strong expression; thinner lines: reduced expression in ΔB1B3 embryos); orange arrowheads indicate the anterior expression stripes. In ΔB1B3 embryos, the posterior domain is strongly reduced or absent, whereas the anterior stripes are comparatively preserved (10 ΔB1B3 embryos analysed). **B:** E9.5. Whole embryos (c, d) and higher-magnification lateral (s–x) and dorsal (y–ad) views of the posterior PSM/tailbud. Snai1 signal is strongly reduced in ΔB1B3 embryos relative to wt, with faint residual expression detectable in most mutants (9 ΔB1B3 embryos analysed). **C:** E10.5. Whole embryos (e, f) and higher-magnification lateral (ae–aj) and dorsal (ak–ap) views of the posterior PSM/tailbud. Distinct spatial patterns of Snai1 expression are observed among embryos of both genotypes, indicating preservation of oscillatory phase variation despite the marked reduction in signal intensity in ΔB1B3 embryos (3 ΔB1B3 embryos analysed). Non-PSM expression domains, including cranial mesenchyme, branchial arches and limb buds, were largely preserved in ΔB1B3 embryos at all stages. Representative embryos are shown.

At E9.5 and E10.5, wild-type *Snai1* PSM expression showed the oscillatory pattern described by [11] with embryos displaying distinct phase configurations, ranging from a broad posterior domain with a prominent anterior stripe to patterns where the anterior component was absent (Fig. 3B, s–u, y–aa; Fig. 3C, ae–ag, ak–am), consistent with the dynamic expression characteristic of the segmentation clock. In ΔB1B3 mutants, oscillatory behaviour was retained at these stages, as evidenced by the presence of distinct phase configurations across embryos (Fig. 3B, v–x, ab–ad; Fig. 3C, ah–aj, an–ap), but overall PSM signal intensity was dramatically reduced. At E9.5 this reduction was near-complete, with only faint residual signal detectable in most mutant embryos (Fig. 3B, d, v–x, ab–ad; n=9). The observation that expression in the cranial mesenchyme, branchial arches and limb buds remained largely unaffected (Fig. 3A, a, b; 3B, c, d; 3C, e, f) indicates that the effect of the ΔB1B3 deletion on *Snai1* expression is specific for the PSM. At E10.5, PSM signal was also reduced relative to wild-type controls, though the difference was less pronounced than at earlier stages (Fig. 3C, f, ah–aj, an–ap; n=3). The retention of oscillatory phase variation in mutants despite reduced signal levels suggests that SCDE controls the amplitude of *Snai1* oscillatory expression in the PSM rather than the oscillatory mechanism itself. *Snai1* expression in the PSM of ΔSCDE mutants, assessed by whole-mount *in situ* hybridization, showed patterns similar to those observed in ΔB1B3 homozygous embryos across the stages examined (Supplementary Fig. 3A, E8.5, n=8; Supplementary Fig. 3B, E10.5, n=3), confirming the relevance of the B1 and B3 blocks for SCDE activity revealed in the transgenic reporter analyses.

To determine whether the enhancer deletions broadly perturbed PSM identity or segmentation clock dynamics, we examined expression of *Lfng*, a core component of the Notch segmentation clock, in ΔB1B3 mutants and wild-type littermates. In both genotypes, *Lfng* displayed the characteristic dynamic stripe patterns in the PSM, with embryos representing distinct phase configurations consistent with ongoing oscillatory activity (Fig. 4A, E8.5, n=10; Fig. 4B, E10.5, n=5). The normal oscillatory behaviour of *Lfng* in mutant embryos demonstrates that the segmentation clock mechanism remains intact in the absence of SCDE and that the reduction in *Snai1* PSM expression observed in these animals does not reflect a general perturbation of clock function. Together, these results establish that SCDE strongly contributes to the level of Snai1 expression in the PSM without affecting the underlying oscillatory machinery.

**Figure 4.**
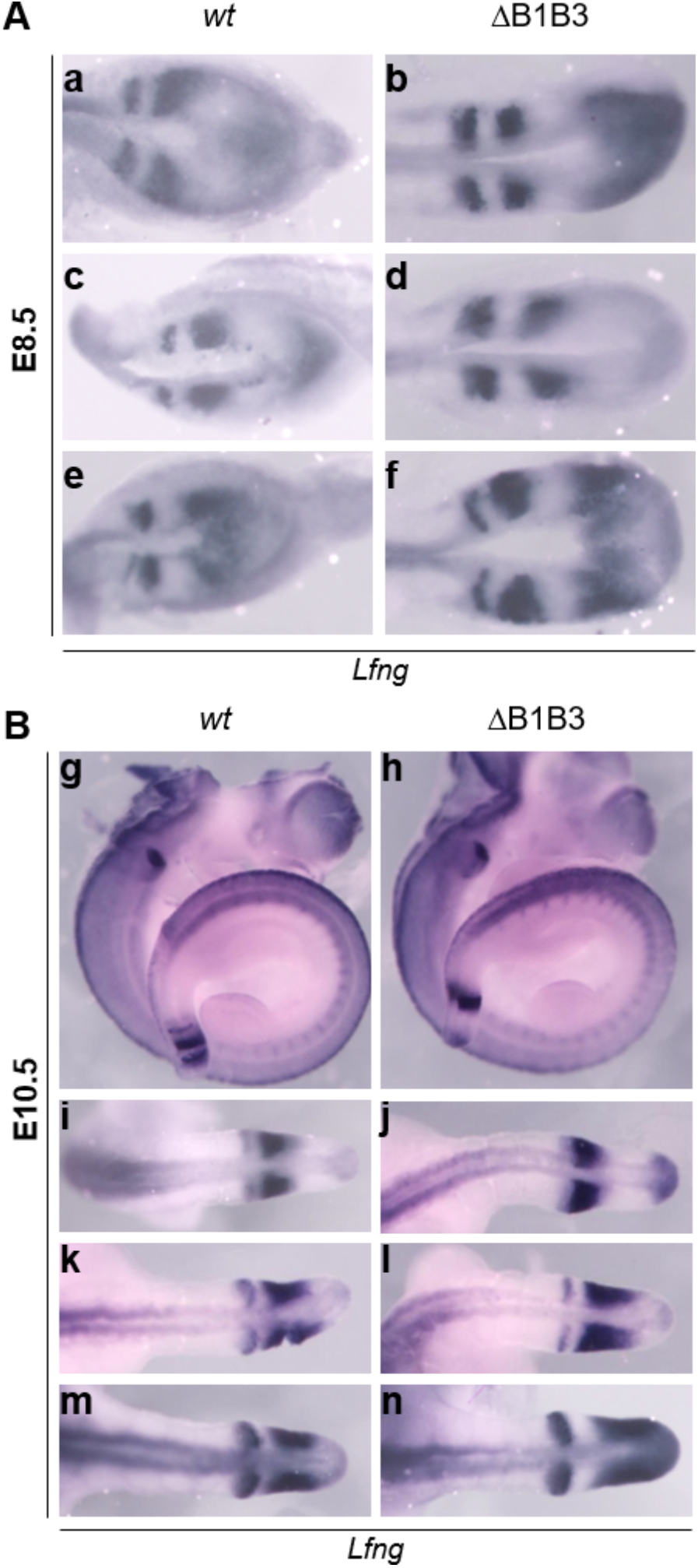
*Lfng* oscillatory expression is preserved in ΔB1B3 mutants. **A:** Whole-mount *in situ* hybridisation for *Lfng* in wild-type littermates and ΔB1B3 embryos at E8.5 (n=10 ΔB1B3 embryos analysed), showing the posterior PSM/tailbud in dorsal view. Representative embryos displaying distinct expression patterns are shown (a–f). **B:** Whole-mount *in situ* hybridisation for *Lfng* in wild-type littermates and ΔB1B3 embryos at E10.5 (n=5 ΔB1B3 embryos analysed), showing whole embryos (g, h) and higher-magnification dorsal views of the posterior PSM/tailbud (i–n). Distinct *Lfng* stripe configurations are observed in both genotypes, consistent with ongoing oscillatory expression and preservation of segmentation clock activity in ΔB1B3 mutants.

### 3.3 T-Cre mediated loss of Snai1 results in perinatal lethality and variable prenatal growth restriction

Although enhancer deletion markedly reduces *Snai1* PSM expression, residual transcription persists in the caudal PSM, particularly at E10.5, precluding a definitive assessment of *Snai1* function in this tissue. To achieve complete elimination of *Snai1* in the PSM and all posterior mesodermal derivatives, we adopted a *Cre/loxP* conditional knockout strategy using a *T-Cre* driver, an approach previously employed to study gene function in posterior mesoderm [29]. We independently generated a T-Cre transgenic line similar to that described by [29] (Supplementary Fig. 4A) and confirmed equivalent recombination activity by ROSA26-lacZ-R reporter analysis (Supplementary Fig. 4B), showing signal in the PSM, tailbud progenitors, somites, and extraembryonic mesoderm (Supplementary Fig. 4C–I). We used this line to obtain conditional mutants in which *Snai1* was eliminated from all *T-Cre* positive territories (*Snai1*^−/flox^; T-Cre, hereafter c*Snai1*).

c*Snai1* mutants were recovered at the expected Mendelian frequency at E9.5–E11.5 (25%, n=32/128), indicating early viability. However, mutant recovery declined substantially by E17.5–E18.5 (14%, n=10/72), and no live mutants were recovered at weaning across 13 litters examined (n=57 pups; Fig. 5A). Two mutants were found dead at P1 and P3 from independent litters, representing the only postnatal animals identified (Fig. 5A; Supplementary Fig. 6v, w). Examination of mutant embryos revealed variable phenotypic severity (Fig. 5B, c; 5D, h), likely reflecting variability in the extent of T-Cre recombination in extraembryonic territories. A subset of mutants presented overt growth retardation, pallor and signs of vascular defects, a phenotype observed with increasing frequency from E11.5 onwards (Fig. 5C, e, 5D, h). The remaining mutants appeared morphologically normal, with some showing subtle growth retardation relative to littermate controls. Strikingly, a proportion of mutants reached E17.5–E18.5 with apparently normal size, gross morphology and visible vasculature, indistinguishable from wild-type littermates by external examination (Fig. 5D, g). The T-Cre line drives recombination in the extraembryonic mesoderm, in addition to the PSM (Supplementary Fig. 4E-I), and variable recombination efficiency in this territory likely underlies the range of prenatal severity observed, consistent with a vascular basis for the lethality, as we discuss below.

**Figure 5.**
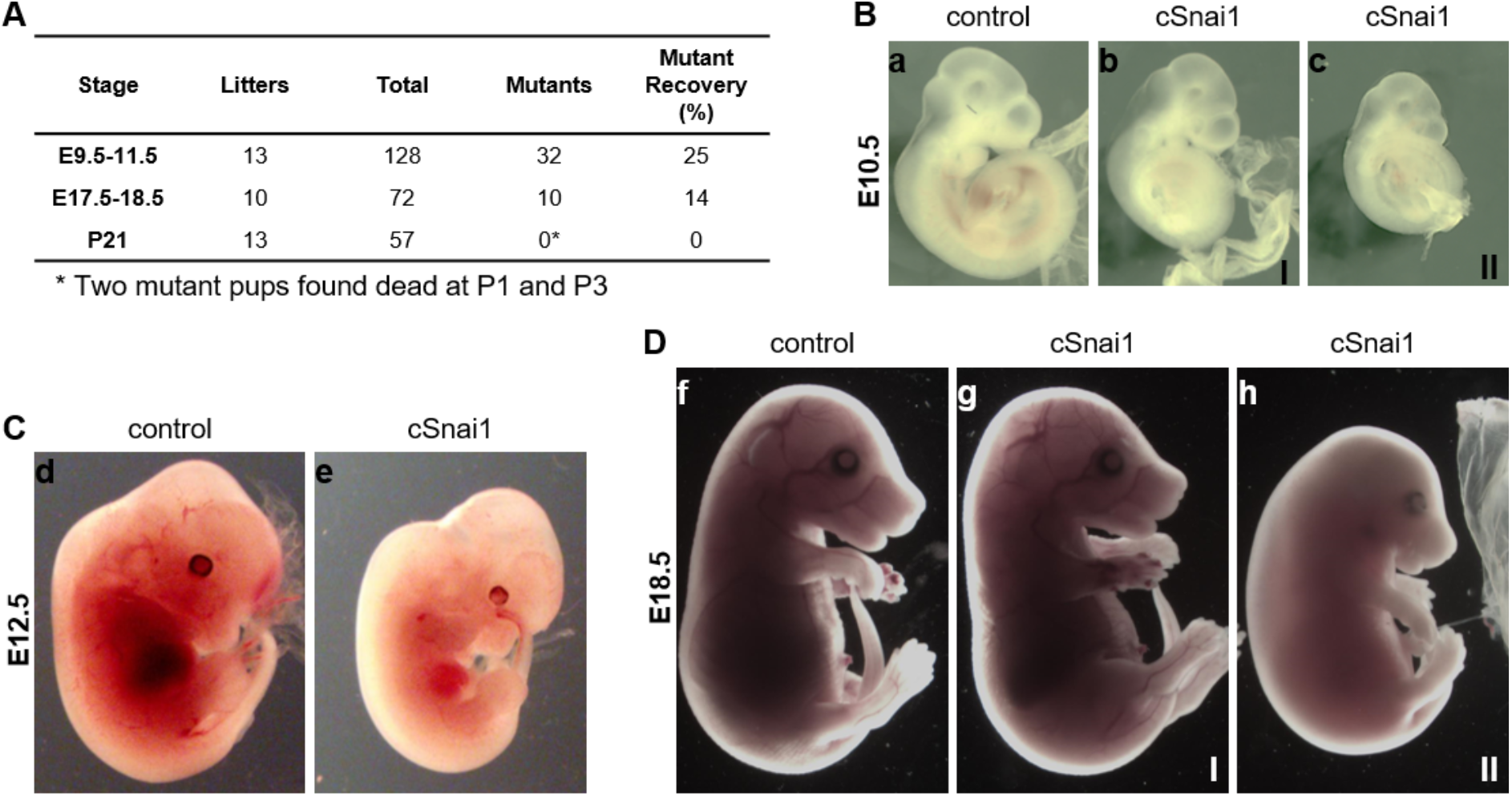
Perinatal lethality and variable phenotypic severity in cSnai1 mutants. **A:** Recovery of cSnai1 mutant embryos and pups at the indicated developmental stages, showing the number of litters examined, total specimens recovered, number of *cSnai1* mutants identified, and mutant recovery as a percentage of total specimens. Mutant recovery decreased from 25% at E9.5–E11.5 to 14% at E17.5–E18.5, and no live mutants were recovered at P21. Two mutant pups were found dead at P1 and P3 (*). **B** Representative control and two *cSnai1* littermates at E10.5, illustrating variable phenotypic severity among mutant littermates (I and II). **C:** Representative control and *cSnai1* embryos at E12.5, illustrating reduced size, pallor and signs of vascular defects in the mutant embryo. **D:** Representative control and *cSnai1* littermates at E18.5, illustrating variable phenotypic severity. Mutant littermate II displays pallor consistent with the vascular phenotype observed at earlier stages.

To assess the functional consequences of *Snai1* loss specifically in the PSM, we focused on the subset of morphologically normal c*Snai1* embryos, in which vascular defects is less likely to confound the interpretation of axial phenotypes. Whole-mount *in situ* hybridisation confirmed efficient *Snai1* elimination in the PSM of these embryos from E9.5 to E11.5 (Fig. 6A, E9.5, n=4; Fig. 6B, E10.5, n=5; Fig. 6C, E11.5, n=2), including the caudal domains where residual expression persisted in SCDE-deletion mutants (Fig. 6A, b, d; 6B, f, i; 6C, n, p). Expression in non-recombined territories, including the cranial mesenchyme and branchial arches, was preserved at all stages examined (Fig. 6A–C), confirming the tissue-specificity of the conditional deletion. Despite complete loss of *Snai1* in the PSM, somite formation and overall embryo morphology appeared grossly normal in these animals, although mutant embryos were frequently smaller than littermate controls even within these morphologically normal mutants (Fig. 6B, g, j), suggesting a subtle but pervasive growth restriction that likely reflects partial vascular defects rather than a consequence of PSM *Snai1* loss.

**Figure 6.**
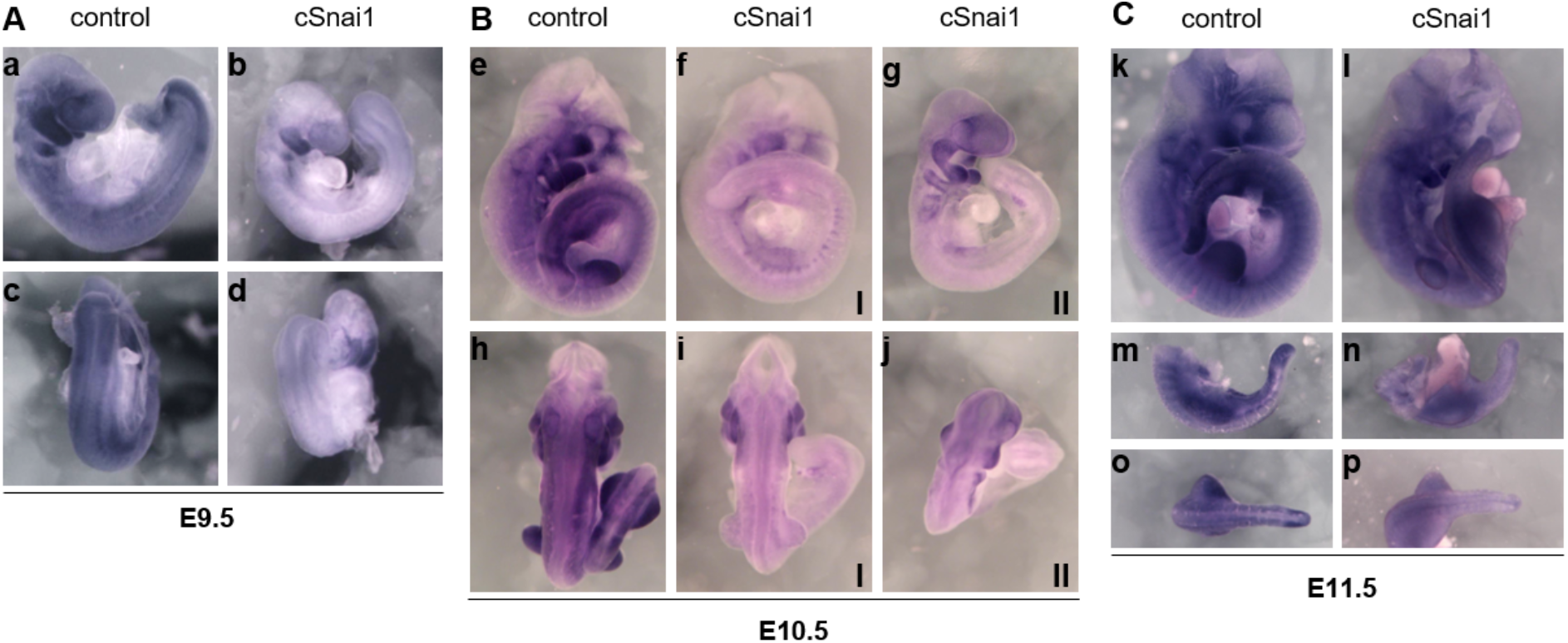
*Snai1* expression is undetectable in the PSM of morphologically unaffected *cSnai1* embryos, with preserved expression in non-recombined territories. **A:** Whole-mount *in situ* hybridisation for *Snai1* in control and morphologically unaffected *cSnai1* embryos at E9.5 (n=4), shown in lateral (a, b) and dorsal (c, d) views. **B:** Whole-mount *in situ* hybridisation at E10.5 (n=5), shown in lateral (e–g) and dorsal (h–j) views; two independent *cSnai1* littermates (I, II) illustrate phenotypic variability, ranging from grossly normal to visibly smaller than control littermates. **C:** Whole-mount *in situ* hybridisation at E11.5 (n=2), shown in lateral view of whole embryos (k, l) and lateral (m, n) and dorsal (o, p) views of the posterior tail region; the cSnai1 embryo shown does not appear visibly smaller than the control. *Snai1* expression is undetectable in the PSM and tailbud of *cSnai1* embryos at all stages examined, while expression in non-recombined territories, including the cranial mesenchyme and branchial arches, is preserved.

### 3.4 Complete loss of Snai1 in the PSM Results in a Fully Penetrant Gain of one Pre-sacral Vertebra

Having confirmed efficient *Snai1* elimination and established that gross embryonic morphology is preserved in morphologically unaffected c*Snai1* embryos, we next assessed whether loss of *Snai1* in the PSM affects somite formation, compartmentalisation or segmentation clock function. For that, we examined a panel of molecular markers in c*Snai1* embryos. *Lfng* expression showed normal oscillatory behaviour in E9.5 c*Snai1* embryos, with distinct phase configurations observed across mutant embryos consistent with ongoing segmentation clock activity (Supplementary Fig. 5A, a–h; n=6). Somite polarity, assessed by *Uncx4.1* expression in the posterior somite compartment at E10, was preserved in all *cSnai1* embryos analysed (Supplementary Fig. 5B, i–p; n=3), with *Uncx4.1* restricted to the posterior half of each somite, indicating that anteroposterior somite polarity is preserved in the absence of *Snai1*. Expression of *Pax1*, a marker of ventral sclerotome, was similarly normal at E10.5 and E12.5, indicating that sclerotome specification is unaffected (Supplementary Fig. 5C, q–t; n=4). Finally, *Tbxt*, marking the notochord and tailbud, was also indistinguishable from controls at E11.5, confirming integrity of axial progenitor territories (Supplementary Fig. 5D, u–z; n=2). Together, these results demonstrate that somite formation, anteroposterior compartmentalisation, sclerotome specification and segmentation clock function are all preserved upon complete loss of *Snai1* in the PSM.

Skeletal analysis of c*Snai1* mutants surviving to E18.5 and the two postnatally recovered animals (Supplementary Fig. 6v, w) revealed a fully penetrant gain of one pre-sacral vertebra in all *cSnai1* mutants for which vertebral formula could be determined (n=8/8; Fig. 7A, B). Littermate controls invariably displayed the expected vertebral formula of 7 cervical, 13 thoracic, 6 lumbar and 4 sacral vertebrae (C7T13L6S4; n=7/7; Fig. 7A, a; 7B). In contrast, all c*Snai1* mutants showed either an additional lumbar vertebra (C7T13L7S4, n=7/8; Fig. 7A, c) or an additional thoracic vertebra (C7T14L6S4, n=1/8; Fig. 7A, b). Sacral vertebral number remained unchanged in all specimens examined. Thus, the total number of pre-sacral vertebrae was increased from 19 in controls to 20 in all mutants, indicating a gain of one axial segment. This phenotype was observed across the full range of morphological severity, including the two postnatally recovered animals (Supplementary Fig. 6v, w; Supplementary Table 2), confirming that it is independent of the vascular defects.

**Figure 7.**
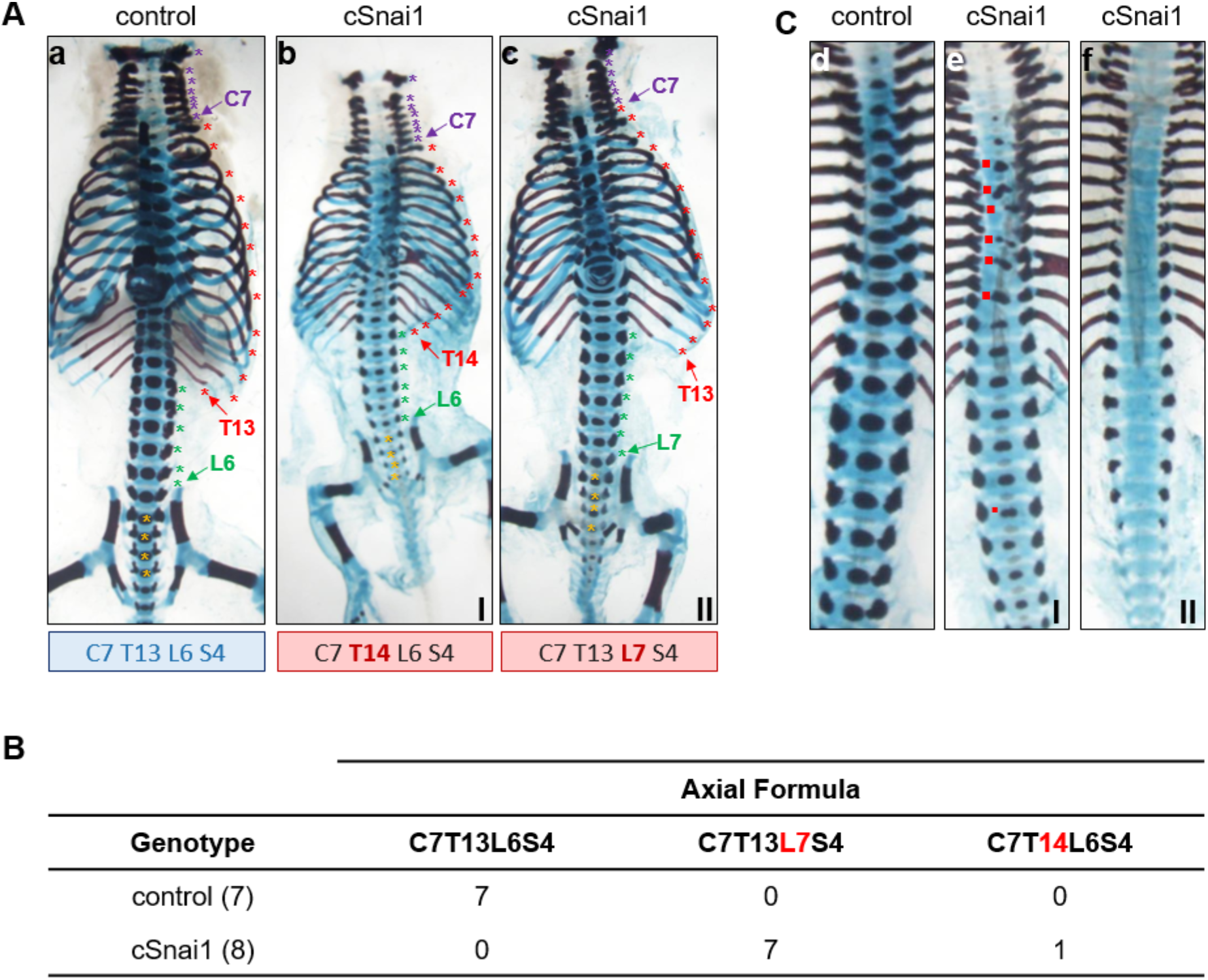
*cSnai1* mutants display a fully penetrant gain of one pre-sacral vertebra. **A:** Alcian Blue/Alizarin Red-stained skeletal preparations of control and cSnai1 littermates (I, II), with vertebrae counted and colour-coded by segment: cervical (C7, purple), thoracic (T13/T14, red), lumbar (L6/L7, green) and sacral (S4, yellow). Vertebral formulae are indicated below each specimen, with the segment differing from control highlighted in red. Control embryos show the expected C7T13L6S4 formula; *cSnai1* littermates show either an additional lumbar vertebra (C7T13L7S4) or, less frequently, an additional thoracic vertebra (C7T14L6S4). **B:** Quantification of vertebral formulae in control (n=7) and cSnai1 mutants for which the vertebral formula could be reliably determined (n=8). All eight assessable *cSnai1* mutants displayed 20 pre-sacral vertebrae, compared with 19 in controls, corresponding to a gain of one pre-sacral vertebra. **C:** Higher-magnification views of the thoracic/lumbar vertebral column in control (d) and two *cSnai1* littermates (e, f). Red squares in (e) indicate focal abnormalities in vertebral ossification centre morphology. Littermate II (f) illustrates the more general reduction in skeletal ossification, affecting both cranial bones and vertebral centres, observed in a subset of growth-restricted mutant embryos.

In addition to the fully penetrant vertebral phenotype, four of the nine mutant specimens analysed (#1, #2, #5 and #6) displayed focal abnormalities affecting vertebral ossification centre morphology within thoracic and lumbar regions (Fig. 7C, e; Supplementary Table2). These alterations were variable in severity and distribution and were not associated with detectable abnormalities in *Uncx4.1* or *Pax1* expression (Supplementary Fig. 5B, C). Additional low-penetrance defects included unilateral proximal rib fusion (n=1/8; Supplementary Fig. 6b), a rudimentary unilateral rib (n=1/8; Supplementary Fig. 6j), and unilateral atlas-occipital fusion (n=1/8; Supplementary Fig. 6c). A subset of growth-restricted embryos displayed a more general reduction in skeletal ossification affecting cranial bones and vertebral centres, which in some cases precluded detailed assessment of vertebral ossification centre morphology (Fig. 7C, f; Supplementary Fig. 6e, g). The most severely affected specimen exhibited marked growth restriction and profound reduction of skeletal ossification, preventing reliable determination of vertebral formula (Supplementary Fig. 6u). Given the variability of these additional skeletal abnormalities and the known activity of *T-Cre* outside the PSM, their relationship to the primary axial phenotype remains unclear and was not investigated further.

Together, these results demonstrate that *Snai1* is dispensable for somite formation, segmentation clock function and somite compartmentalisation, but contributes to the precise determination of axial segment number, with its loss consistently resulting in the formation of one additional pre-sacral vertebra. In addition, the subtle phenotypes observed in the vertebrae indicate that *Snai1* might play a role in later morphogenetic processes in somite derivatives.

## 4. Discussion

The results presented here identify SCDE as a bona fide *cis*-regulatory enhancer involved in the control of *Snai1* expression. Although transgenic reporter assays indicate that SCDE might contribute to *Snai1* expression in various embryonic regions, deletion of this enhancer in mouse embryos indicates that its most relevant activity lies in regulating *Snai1* expression in the PSM. This implies the existence of additional, as yet unidentified, regulatory elements that control *Snai1* expression in other domains, such as the cranial mesoderm or branchial arches. Functional dissection of SCDE using transgenic reporter assays revealed that it comprises two core regulatory modules, B1 and B3, which act redundantly to drive expression in the PSM and tailbud, while contributing differently to craniofacial and limb expression domains. Redundancy between these two modules is supported by the observation that individual deletion of either B1 or B3 alone preserves most axial enhancer activity, whereas simultaneous deletion abolishes expression almost completely. Functional redundancy has been widely observed for developmental enhancers controlling genes with complex spatiotemporal expression patterns [30,31], where partially redundant *cis*-regulatory modules confer robustness while simultaneously allowing tissue-specific specialisation.

SCDE has been shown to be a direct transcriptional target of Msgn1, a master regulator of paraxial mesoderm differentiation, as indicated by ChIP analyses in mouse tissues and further supported by in vitro data [19]. Our work provides direct functional evidence that SCDE might indeed be involved in Msgn1-dependent activation of *Snai1* in the PSM. Notably, *Snai1* expression is absent from the PSM of *Msgn1*-null embryos but remains detectable in non-PSM territories [19]. This pattern resembles that of SCDE-deleted mutants, which is consistent with SCDE constituting a major *cis*-regulatory interface through which Msgn1 activates *Snai1* during axis elongation. Nevertheless, residual PSM expression persists in ΔB1B3 and ΔSCDE lines, indicating that additional *cis*-regulatory elements elsewhere in the *Snai1* locus contribute to its regulation by Msgn1 in the PSM. This residual activity is spatially and temporally patterned: it is most prominent in the anterior PSM at E8.5 and becomes more significant at E10.5. Notably, oscillatory *Snai1* activity was still observed in ΔB1B3 and ΔSCDE mutant embryos despite the markedly reduced expression levels in the PSM, suggesting that the remaining activity is supported by additional regulatory elements. Together, these observations support a model in which SCDE functions as a regulatory element controlling posterior *Snai1* expression during early axis elongation. However, its primary role appears to be defining the overall *Snai1* expression levels within the PSM, thereby delimiting the range within which other regulatory elements, especially those responsible for its oscillatory expression, can subsequently operate.

The apparent existence of dedicated regulatory mechanisms controlling oscillatory *Snai1* expression in the PSM contrasts with the observation that *Snai1* expression is dispensable for the segmentation process, as somitogenesis proceeds normally in embryos lacking *Snai1* in the PSM. This raises an important conceptual question: why does *Snai1* exhibit dynamic expression in the PSM [11]? Clearly, *Snai1* is not involved in regulating the dynamic expression of other oscillatory genes, as such a role would lead to abnormal somitogenesis in the *Snai1* mutants. Consistent with this, the oscillatory behaviour of *Lfng* appears unaffected in embryos lacking *Snai1* in the PSM. A possible explanation is that oscillatory *Snai1* expression simply reflects the dynamic regulatory landscape of the PSM. In this scenario oscillating factors controlling the segmentation clock also act on *Snai1* enhancers that might be involved in the regulation of this gene in other embryonic areas that are also accessible in the PSM. *Snai1* would thus represent another of the genes showing oscillatory expression in the PSM without direct contribution to oscillator function [32–34]. Notably, while oscillatory *Snai1* expression at E9.5 and later stages was reproducible, we did not detect obvious dynamic expression at E8.5, as all nine embryos examined displayed similar patterns.

Inactivation of *Snai1* in the posterior mesoderm did not disrupt somite formation; however, it may still contribute to sclerotome morphogenesis. In the limited number of embryos recovered at stages suitable for skeletal analysis, we observed vertebral body abnormalities of variable penetrance, ranging from subtle defects to an apparent absence of ossified vertebral bodies. Although the latter phenotype may partially reflect a developmental delay, the overall spectrum of defects may reflect alterations in later sclerotome maturation or vertebral morphogenesis. These findings contrast sharply with the complete absence of axial skeleton reported upon *Msgn1* inactivation [35], indicating that *Snai1* plays a minor, if any, role downstream of *Msgn1*, and that the vertebral phenotype is unlikely to arise from a requirement for *Snai1* in the PSM. Because T-Cre–mediated recombination includes somitic expression domains of *Snai1*, the most plausible interpretation is that the vertebral defects in these conditional mutants reflect a function of *Snai1* during later morphogenetic events of sclerotome differentiation. Consistent with this view, somite patterning and early sclerotome induction appear largely preserved in the conditional mutants, supporting the conclusion that *Snai1* is required at a subsequent stage of vertebral development rather than during initial somite formation.

*Snai1* conditional inactivation using the T-Cre driver resulted in a fully penetrant gain of one pre-sacral vertebra in all mutants examined. The position of the sacrum reflects the transition from trunk to tail development, a developmental shift that is functionally connected with the relocation of the neuromesodermal progenitor (NMP) population [36,37] from the epiblast to the tail bud. This relocation is regulated by a *Tgfbr1*/*Snai1* network that induces an incomplete EMT in these progenitors as previously described [21]. Reporter analysis using the ROSA26-lacZ-R line (Supplementary Fig. 4), is consistent with T-Cre activity in at least a subset of NMPs, as evidenced by labelling within the neural tube (Supplementary Fig. 4D-F). Consequently, the additional pre-sacral vertebra observed in *cSnai1* mutants may reflect a slight delay in the initiation of NMP relocation to the tail bud. The pattern of neural tube labelling suggests that *Snai1* inactivation within NMPs is partial and increases progressively during development. This graded inactivation would explain why the trunk-to-tail transition is not completely disrupted, in contrast to the phenotype resulting from early *Snai1* deletion in the epiblast, which removes *Snai1* from all embryonic lineages, including the entire NMP population. This model remains hypothetical, as NMP number and relocation dynamics were not directly assessed in the present study.

The axial patterning phenotype described above is accompanied by perinatal lethality in a subset of c*Snai1* mutants, a finding that likely reflects *Snai1* function in the extraembryonic mesoderm and/or vascular mesodermal territories. Indeed, the T-Cre line drives recombination in the embryonic posterior and extraembryonic mesoderm, in addition to the PSM, affecting both endothelial and supporting mesodermal cell populations across embryonic and yolk sac vasculature. Whether the observed lethality reflects a cell-autonomous requirement for *Snai1* in endothelial cells, in the surrounding mesoderm, or both, cannot be determined from the current data. Nevertheless, this phenotype is consistent with the embryonic lethality and vascular defects reported upon epiblast-specific *Snai1* deletion [38], and with the demonstration that endothelial-specific loss of *Snai1* causes early lethality associated with profound defects in vascular remodelling and yolk sac perfusion [39]. These vascular defects may have secondarily affected endochondral ossification, ultimately contributing to the reduced skeletal ossification observed in growth-restricted mutants [40]. This interpretation is further supported by the observation that conditional deletion of *Snai1* alone in the limb mesenchyme does not produce detectable skeletal defects [41]. The full characterisation of this phenotype lies beyond the scope of the present study.

In conclusion, this work identifies SCDE as a *cis*-regulatory enhancer involved in *Snai1* expression in the PSM, although it appears dispensable for the oscillatory component of this expression, which persists even when SCDE activity is severely reduced or abolished. Together with the normal segmentation, somite compartmentalisation and clock gene dynamics observed upon efficient loss of *Snai1* expression in the PSM, our results indicate that neither *Snai1* expression nor its oscillatory activity is required for somitogenesis. Whether this dissociation between cyclic expression and clock requirement extends to other PSM-expressed genes remains an open question.

## CRediT authorship contribution statement

Ana Casaca: Conceptualization, Investigation, Formal analysis, Funding acquisition, Writing – original draft, Writing – review & editing.

Rita Aires: Conceptualization, Investigation, Methodology, Writing – review & editing. Ana Nóvoa: Investigation.

Moisés Mallo: Conceptualization, Supervision, Funding acquisition, Writing – original draft, Writing – review & editing.

## AI usage disclosure

Microsoft Copilot was used solely for language editing to improve grammar and clarity. All changes were reviewed and approved by the authors, who confirm that the scientific content, conclusions, and interpretation of the results remained unchanged.

## Acknowledgements

We would like to thank Angela Nieto for the *Snai1* cDNA. We also thank the members of the Mallo lab for continuous support at different stages of this project and the GIMM Rodent scientific platform for their help with animal housing. This work was funded by Fundação para a Ciência e a Tecnologia (FCT) grants 2023.12793.PEX to AC (DOI: 10.54499/2023.12793.PEX) and 2022.01629.PTDC to MM (DOI: 10.54499/2022.01629.PTDC), and supported by GIMM-CARE (funded by the European Union under grant agreement No. 101060102. GIMM-CARE is co-funded by the Portuguese Government, the Foundation for Science and Technology (FCT), ARICA – Investimentos, Participações e Gestão, Jerónimo Martins, the Gulbenkian Institute for Molecular Medicine, and CAML – Lisbon Academic Medical Centre) [doi.org/10.3030/101060102], and by national funds through FCT under the Associate Laboratory programme (LA/P/0082/2020) [doi.org/10.54499/LA/P/0082/2020], and under the R&D Unit funding programme (UID/06357/2025) [doi.org/10.54499/UID/06357/2025].

## Data Availability

Data will be made available on request

