## Supplementary figures and tables for "Normal Somitogenesis Despite the Absence of *Snai1* in the Presomitic Mesoderm"

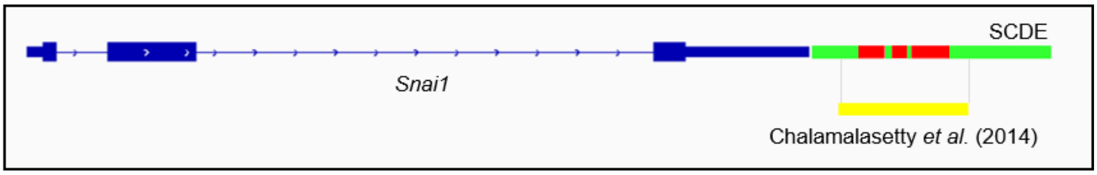


**Supplementary Figure 1. The SCDE encompasses the previously identified Msgn1-responsive *Snai1* enhancer region.** Schematic representation of the *Snai1* locus, showing exons (blue boxes) and the downstream SCDE region (green), subdivided into conserved blocks B1, B2 and B3 (red). The genomic interval previously identified as a Msgn1-responsive element by Chalamalasetty et al. (2014) is indicated in yellow, illustrating that it corresponds to a subregion within the broader SCDE identified here.

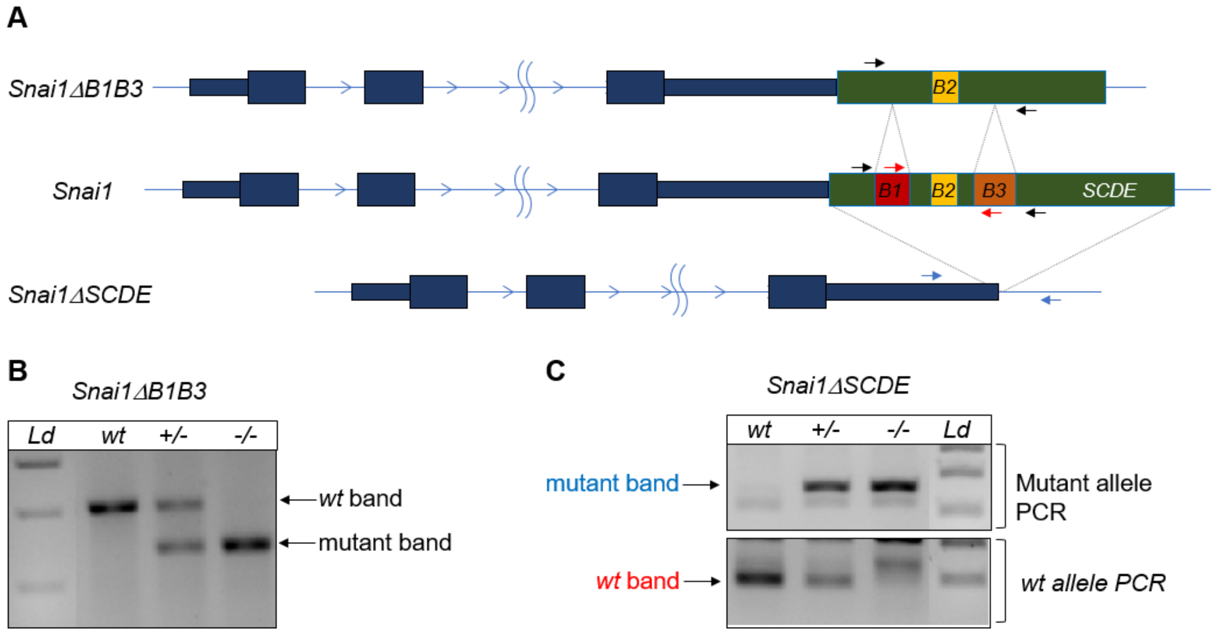


**Supplementary Figure 2. Generation and validation of Snai1ΔB1B3 and Snai1ΔSCDE mouse lines.**

**A:** Schematic representation of the *Snai1* locus and the two CRISPR/Cas9-generated deletion alleles. The wild-type *Snai1* locus (middle) shows exons (dark blue boxes) and the downstream SCDE region (green), with conserved Blocks B1 (red), B2 (yellow) and B3 (orange) indicated. The Snai1ΔB1B3 allele (top) lacks Blocks B1 and B3 while retaining Block B2 and the remaining SCDE sequence. The Snai1ΔSCDE allele (bottom) lacks the entire SCDE region. Arrows indicate the primers used for genotyping: black arrows, primer pair flanking Blocks B1 and B3, used for the Snai1ΔB1B3 line and amplifying the wild-type (473 bp) and mutant (315 bp) alleles; blue arrows, primer pair flanking the deletion breakpoint in the Snai1ΔSCDE allele, amplifying the mutant allele (361 bp); red arrows, primer pair located within Blocks B1 (forward) and B3 (reverse), amplifying the wild-type allele of the Snai1ΔSCDE line (253 bp). **B:** Representative PCR genotyping gel for the Snai1ΔB1B3 line, showing wild-type (wt), heterozygous (+/−) and homozygous mutant (−/−) genotypes, distinguished by wt and mutant band sizes. **C:** Representative PCR genotyping gels for the Snai1ΔSCDE line, showing mutant allele PCR (top gel) and wild-type allele PCR (bottom gel) across wt, +/− and −/− genotypes. Ld: DNA ladder

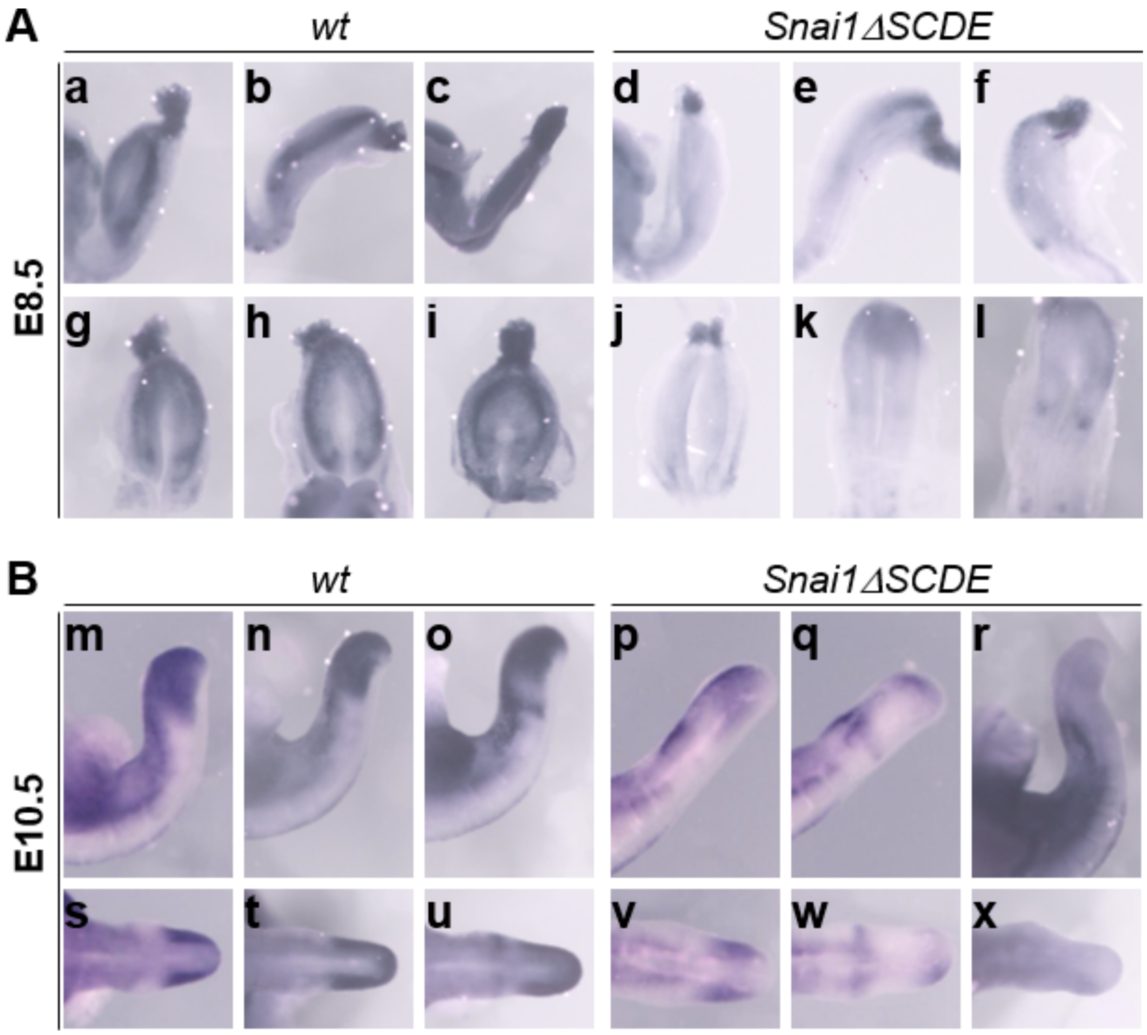


**Supplementary Figure 3. *Snai1* expression is reduced in the PSM of ΔSCDE mutant embryos.**

**A:** Whole-mount *in situ* hybridisation for *Snai1* in control and ΔSCDE homozygous embryos at E8.5 (n=8), shown in lateral view of the tail for control (a–c) and ΔSCDE (d–f) embryos, and in dorsal view of the posterior tail region for control (g–i) and ΔSCDE (j–l) embryos. **B:** Whole-mount *in situ* hybridisation at E10.5 (n=3), shown in lateral view of the tail for control (m–o) and ΔSCDE (p–r) embryos, and in dorsal view for control (s–u) and ΔSCDE (v–x) embryos. At both stages, ΔSCDE embryos show markedly reduced *Snai1* signal in the PSM compared to controls, while retaining detectable expression, consistent with the phenotype observed in Snai1ΔB1B3 mutants (Figure 3).

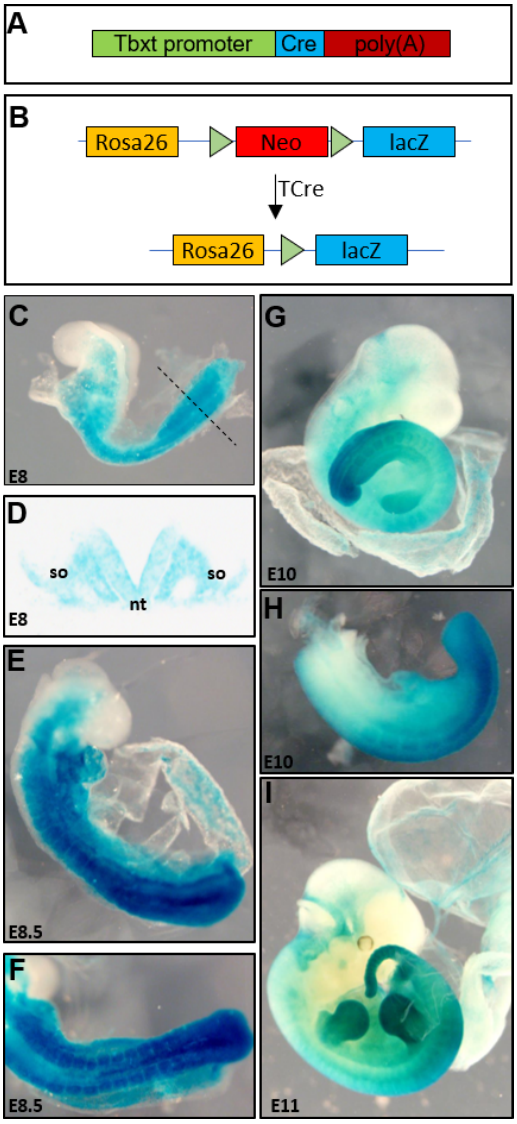


**Supplementary Figure 4. Generation of the T-Cre:ROSA26-lacZ reporter line and validation of T-Cre recombination activity.**
**A:** Schematic representation of the T-Cre transgene, comprising Cre recombinase under the control of the *Tbxt* primitive streak-specific promoter followed by a poly(A) signal. **B:** Schematic representation of the ROSA26-lacZ-R reporter locus before and after Cre-mediated recombination. Cre-mediated excision of the loxP-flanked Neo cassette activates lacZ expression from the endogenous ROSA26 locus. **C:** E8 embryo showing recombination in the primitive streak and posterior mesoderm; the plane of section shown in D is indicated by the dashed line. **D:** Transverse section through the level indicated in C, showing β-galactosidase activity in somites (so) and neural tube (nt). **E, F:** E8.5 embryos showing extensive recombination in posterior and paraxial mesoderm/somites in a lateral (E) and dorsal (F) view, with additional staining in the yolk sac and neural tube. **G:** E10 embryo showing recombination in posterior embryonic and extraembryonic tissues, including the yolk sac and limb buds. **H:** Higher-magnification lateral view of the posterior region shown in G. **I:** E11 embryo showing recombination in posterior embryonic tissues, limb buds and extraembryonic membranes.

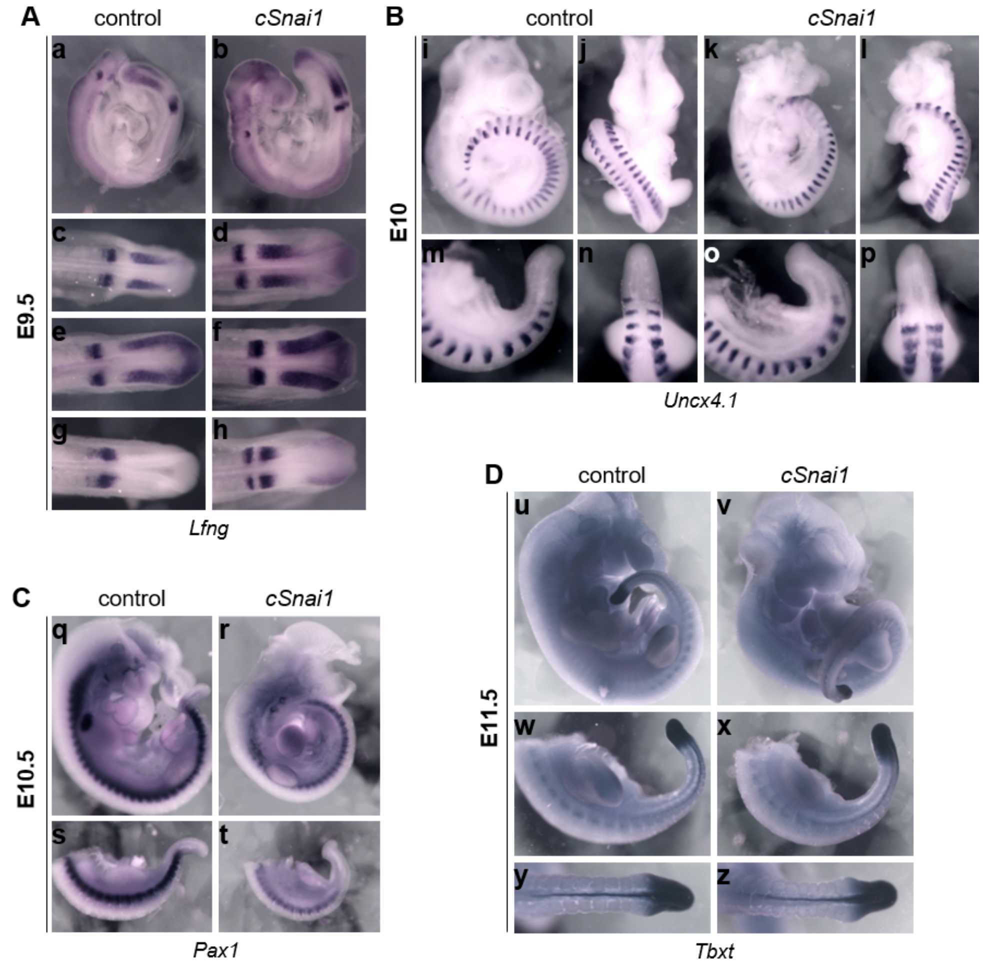


**Supplementary Figure 5. Segmentation clock and somite patterning are preserved in *cSnai1* mutants.**

Whole-mount *in situ* hybridisation for segmentation clock and somite patterning markers in control and *cSnai1* embryos. **A:** *Lfng* expression in representative control (a) and *cSnai1* (b) embryos at E9.5 (6 *cSnai1* embryos analysed). Dorsal views of the posterior PSM of three control (c, e, g) and three cSnai1 (d, f, h) embryos show distinct phase configurations in both genotypes, indicating normal oscillatory clock activity. **B:** *Uncx4.1* expression at E10 in a representative control (i, j, m, n) and cSnai1 (k, l, o, p) embryo, shown in lateral (i, k) and dorsal (j, l) views, with higher-magnification lateral (m, o) and dorsal (n, p) views of the tail. *Uncx4.1* is restricted to the posterior half of each somite in both genotypes, indicating that somite polarity is preserved (3 *cSnai1* embryos analysed). **C:** *Pax1* expression, marking the ventral sclerotome, in a representative control and *cSnai1* embryo at E10.5, shown at low magnification (q, r) and in lateral view of the posterior tail (s, t). Sclerotome specification is unaffected in *cSnai1* mutants (4 cSnai1 embryos analysed: two at E10.5 and two at E12.5). **D:** *Tbxt* expression, marking the notochord and tailbud, in a representative control and *cSnai1* embryo at E11.5, shown at low magnification (u, v) and in lateral (w, x) and dorsal (y, z) views of the tail. Notochord and tailbud integrity are preserved in *cSnai1* mutants (2 *cSnai1* embryos analysed)

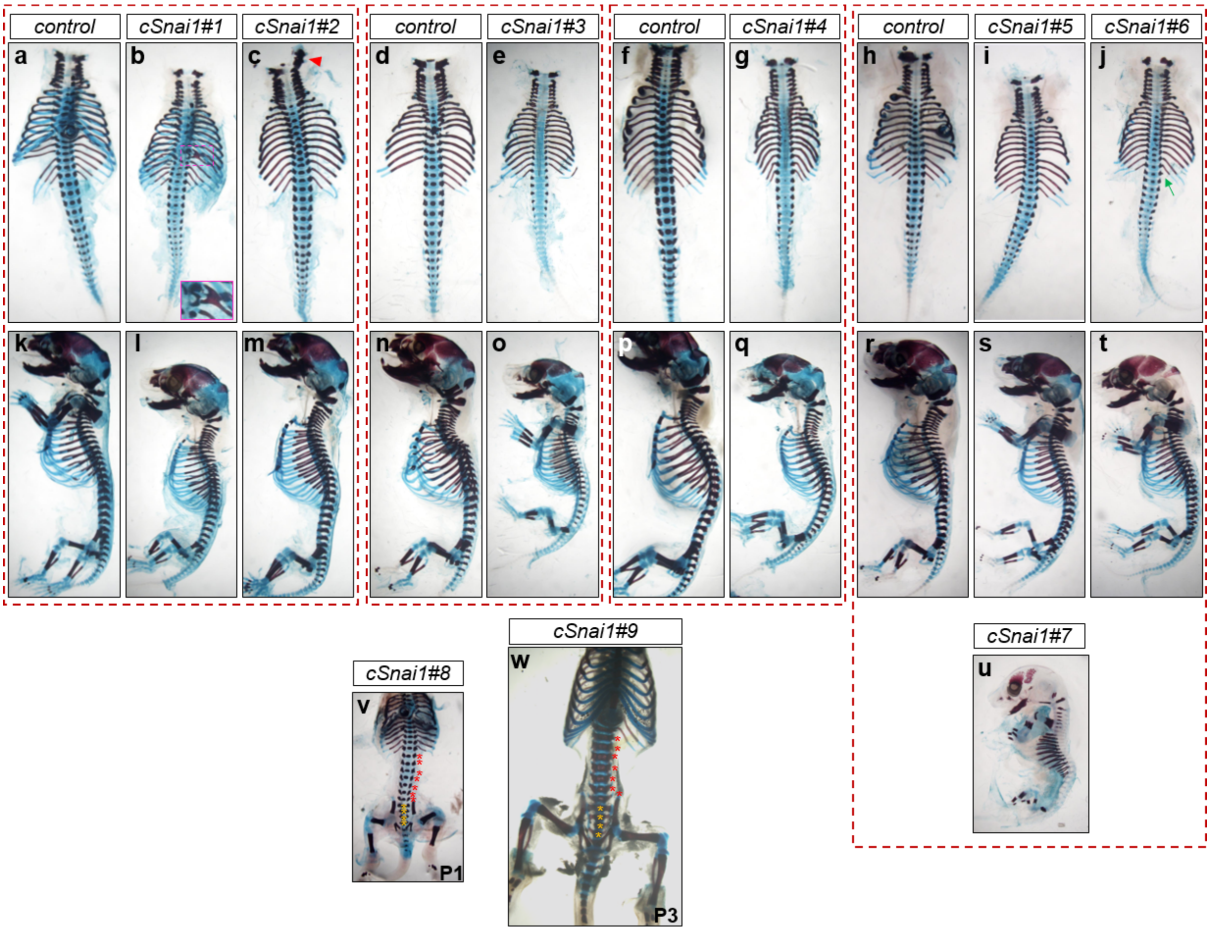


**Supplementary Figure 6. Skeletal analysis of cSnai1 mutant embryos and neonates.** Alcian Blue/Alizarin Red-stained skeletal preparations of control and *cSnai1* mutant specimens. Dashed red boxes group littermates processed together. Litter 1 (a–c, k–m): control, cSnai1#1 and cSnai1#2, shown in dorsal thoracolumbar view (a–c) and lateral whole-skeleton view (k–m). The magenta box in (b) indicates the region of unilateral proximal rib fusion in cSnai1#1, shown magnified in the inset. The red arrowhead in (c) indicates the site of unilateral atlas-occipital fusion in cSnai1#2. (d–e, n–o) Litter 2: control and cSnai1#3. (f–g, p–q) Litter 3: control and cSnai1#4. (h–j, r–t) Litter 4: control, cSnai1#5, cSnai1#6 and cSnai1#7. The green arrow in (j) indicates the rudimentary unilateral rib in cSnai1#6. (u) cSnai1#7, the most severely affected specimen, showing marked growth restriction and reduced skeletal ossification. (v) cSnai1#8, recovered postnatally at P1. (w) cSnai1#9, recovered postnatally at P3. In (v, w), red asterisks indicate lumbar vertebrae and yellow asterisks indicate sacral vertebrae.

Supplementary Table1. Oligonucleotides and probes used in this work.

| Genotyping primers | | |
|---|---|---|
| ΔB1B3<br>Mutant band=325 bp<br>wt band= 427 bp | Forward | GGGGGCACCATAAAGGGACTTCC |
| | Reverse | AGCACCAACTAGCCTGAGGT |
| ΔSCDE<br>Mutant band=361 bp | Forward | TGAACTTTGGGGCCTGGGC |
| | Reverse | AGCCCCTTGGCTGCATTCTGG |
| Snai1ΔSCDE<br>wt band=253 bp | Forward | AATGGGCTGGCTGGAGCCAGG |
| | Reverse | TTGGAGCTGGGCTGGCTGAAGC |
| Snai1- | Forward | TGAAAGCGGCTCTGTTCAGT |
| | Reverse | CGGGCTTAGGTGTTTTCAGAC |
| Tcre | Forward | GGGCAGCTGCTCGGTACTTC |
| | Reverse | TGCGAACCTCATCACTCGTTC |
| Snai1flox<br>Snai1flox band=365bp<br>wt band= 249 bp | Forward | TGAAAGCGGCTCTGTTCAGTG |
| | Reverse | CTGCTGCACCCCTACTATGTG |
| **Cloning of SCDE reporter constructs** | | |
| SCDE | Forward (MluI) | acgcgtGGGGGCACCATAAAGGGACTTCC |
| | Reverse (BglII) | AGATCTCCATCCCTCTTCCTCCACCTCC |
| B1 | Forward (XhoI) | ggctcgagGGGAGGATGTCTTTCCTGTTCCC |
| | Reverse (BglII) | ccagatctGCAGGAAGCTGTATGACCCC |
| B2 | Forward (XhoI) | ggctcgagggtgacctcaactctgagggc |
| | Reverse (BglII) | ccagatctCCTGCTCAGCTCCCCACAGG |
| B3 | Forward (XhoI) | ggctcgagGGAGCTTCAGCCAGCCCAGC |
| | Reverse (BglII) | ccagatctCCTGCTCAGCTCCCCACAGG |
| ΔB1 I | Forward (MluI) | acgcgtGGGGGCACCATAAAGGGACTTCC |
| | Reverse | GGTCCTCCAATTCAGCTGGGAGGGGCAGCTA |
| ΔB1 II | Forward | CCCAGCTGAATTGGAGGACCAGGGCTGGAGC |
| | Reverse (BglII) | AGATCTCCATCCCTCTTCCTCCACCTCC |
| ΔB2 I | Forward (MluI) | acgcgtGGGGGCACCATAAAGGGACTTCC |
| | Reverse | GCCGACGCCTAGCACCAACTAGCCTGAGGT |

| | | |
|---|---|---|
| ∆B2 II | Forward | AGTTGGTGCTAGGCGTCGGCTGGGAGCTTC |
| | Reverse (BglII) | AGATCTCCATCCCTCTTCCTCCACCTCC |
| ∆B3 I | Forward (MluI) | acgcgtGGGGGCACCATAAAGGGACTTCC |
| | Reverse | GGCCTTCAGCCCAGCCGACGCCTGCTCAGC |
| ∆B3 II | Forward | CGTCGGCTGGGCTGAAGGCCTACTGTGTCC |
| | Reverse (BglII) | AGATCTCCATCCCTCTTCCTCCACCTCC |
| **Oligonucleotides for *in situ* hybridization probes** | | |
| Snai1CDS probe | Forward XbaI | ccTCTAGAAtgccgcgctccttcctggtc |
| | Reverse(EcoRI) | ccgaattcTCAGCGAGGGCCTCCGGAGC |
| **CRISPR/Cas9 Reagents** | | |
| Snai1∆B1B3 | gRNA 5’ | TGCCCCTCCCAGCTGAAGGG |
| | gRNA 3’ | GCTGGGCATCAACGAGCTGA |
| | Forward for Donor ssDNA | TGAACTTTGGGGCCTGGGC |
| | Reverse for Donor ssDNA | AGGTGAGAGTCAGGGCCAGG |
| Snai1∆SCDE | gRNA 5’ | TCTGCCCCAAAGACCTGCTA |
| | gRNA 3’ | TGGAGGTGGAGGAAGAGGGA |
| | Donor ssDNA | TGGGAAGGTGCTTTGTGGGGCTTAGCCACAG<br>CCTGGCCAGGAGCTCTGGCTCCACCATAGGA<br>GGGTTGACATTCCAGAGAGGGAGGAGGAGG<br>ATGGGAAGGATGGAGGTGGAGGAAGAGG |

Supplementary Table2. Vertebral formulae and skeletal findings in individual cSnai1 mutant specimens

| Specimen ID | Developmental Stage | Vertebral formula | Other findings |
|---|---|---|---|
| #1 | E18.5 | C7T14L6S4 | Unilateral proximal rib fusion; vertebral ossification centre abnormalities |
| #2 | E18.5 | C7T13L7S4 | Unilateral atlas-occipital fusion; vertebral ossification centre abnormalities |

| | | | |
|---|---|---|---|
| #3 | E18.5 | C7T13L7S4 | General ossification reduction |
| #4 | E18.5 | C7T13L7S4 | General ossification reduction |
| #5 | E18.5 | C7T13L7S4 | Vertebral ossification centre abnormalities |
| #6 | E18.5 | C7T13L7S4 | Rudimentary unilateral rib; vertebral ossification centre abnormalities |
| #7 | E18.5 | Not assessable | Marked growth restriction; Severe ossification reduction |
| #8 | P1 | C7T13L7S4 | — |
| #9 | P3 | C7T13L7S4 | — |

Vertebral formula and additional skeletal findings for each cSnai1 mutant specimen analysed by skeletal preparation, corresponding to the specimens shown in Supplementary Figure 6. Specimen #7could not be reliably assessed for vertebral formula due to severe ossification reduction associated with marked growth restriction (see main text).
